# pGpG-signaling regulates virulence and global transcriptomic targets in *Erwinia amylovora*

**DOI:** 10.1101/2024.01.12.575434

**Authors:** Roshni R. Kharadi, Brian Y. Hsueh, Christopher M. Waters, George W. Sundin

**Affiliations:** Department of Plant, Soil and Microbial Sciences, Michigan State University, East Lansing, Michigan, USA; Department of Microbiology and Molecular Genetics, Michigan State University, East Lansing, Michigan, USA

## Abstract

Cyclic-di-GMP (c-di-GMP) is a critical bacterial second messenger that enables the physiological phase transition in *Erwinia amylovora*, the phytopathogenic bacterium that causes fire blight disease. C-di-GMP generation is dependent on diguanylate cyclase enzymes while the degradation of c-di-GMP can occur through the action of phosphodiesterase (PDE) enzymes that contain an active EAL and/or a HD-GYP domain. The HD-GYP-type PDEs, which are absent in *E. amylovora*, can directly degrade c-di-GMP into two GMP molecules. PDEs that contain an active EAL domain, as found in all active PDEs in *E. amylovora,* degrade c-di-GMP into pGpG. The signaling function of pGpG is not fully understood in bacterial systems. A transcriptomic approach revealed that elevated levels of pGpG in *E. amylovora* impacted several genes involved in metabolic and regulatory functions including several type III secretion and extracellular appendage related genes. The heterologous overexpression of an EAL or HD-GYP-type PDE in different background *E. amylovora* strains with varying c-di-GMP levels revealed that in contrast to the generation of pGpG, the direct breakdown of c-di-GMP into GMP by the HD-GYP-type PDE led to an elevation in amylovoran production and biofilm formation despite a decrease in c-di-GMP levels. The breakdown of c-di-GMP into pGpG (as opposed to GTP) also led to a decrease in virulence in apple shoots. The expression of *hrpS* was significantly increased in response to the breakdown of c-di-GMP into pGpG. Further, our model suggests that a balance in the intracellular ratio of pGpG and c-di-GMP is essential for biofilm regulation in *E. amylovora*.

**Importance:** c-di-GMP is the keystone molecule for regulating the transition from motility to biofilm formation in most bacteria. Interestingly, there are two distinct enzymatic phosphodiesterase (PDE) domains, termed EAL and HD-GYP, that degrade c-di-GMP. EAL domains cleave one bond of the cyclic ring to generate pGpG while HD-GYP enzymes cleave c-di-GMP to two GMP molecules. A central question regarding c-di-GMP signaling is has whether or not pGpG itself functions as a signaling molecule. Here we demonstrate in the plant pathogen *Erwinia amylovora* that pGpG specifically regulates genes and contributes to biofilm formation and disease progression.

## Introduction

*Erwinia amylovora* is the causal agent of the economically important fire blight disease that affects apple and pear production worldwide (1). During the shoot blight infection cycle, *E. amylovora* cells must successfully transition from type III secretion (T3S) driven virulence in the apoplast region to a systemic form of host colonization mediated by the formation of biofilms within the xylem vessels (2). The ubiquitous bacterial second messenger cyclic-di-GMP (c-di-GMP) is a critical signaling factor that enables this tissue specific virulence transition in *E. amylovora* (3, 4). The synthesis of c-di-GMP involves the dimerization of two GTP molecules enzymatically facilitated by diguanylate cycle (DGC) enzymes (5). Phosphodiesterase (PDE) enzymes that contain an active EAL domain can hydrolyze c-di-GMP into 5′-phosphoguanylyl-(3′ → 5′)-guanosine (pGpG), which can be further degraded into GMP by oligoribonucleases (Orn) or pGpG specific PDEs (5–7). In contrast, PDEs with active HD-GYP domains can directly degrade c-di-GMP into GMP (8). *E. amylovora* encodes five active DGCs, three active PDEs (3, 4), and all three active PDEs in *E. amylovora* only contain an EAL domain (4).

In *E. amylovora*, c-di-GMP negatively regulates flagellar motility in vitro (3, 4, 9). Further, T3S is negatively regulated by c-di-GMP through the transcriptional repression of *hrpL* which encodes for an alternative sigma factor (σ^54^) that regulates the transcription of T3S genes (3, 4, 9, 10). This further leads to a reduction in DspE translocation within the leaf apoplast (4). *hrpL* is under the transcriptional regulation of HrpS which is an NtrC family σ^54^-dependent bacterial enhancer binding protein (bEBP) (11–13). HrpS can regulate its own expression through the transcriptional activation of the *hrpL* promoter (12, 13). HrpXY and the Rcs phosphorelay are two component signal transduction systems that also regulate *hrpS* transcription (13, 14). HrpS consists of an N-terminal receiver (Rec) domain, a central AAA+ (ATPase) involved in ATP hydrolysis and σ^54^ interaction, and a C-terminal HTH (helix-turn-helix) DNA binding domain (11, 12, 15). Other bEBPs such as FlrA in *Vibrio cholerae* and FleQ in *Pseudomonas aeruginosa* are inhibited by c-di-GMP reducing bacterial motility (16, 17). Alternatively, the *V. cholerae* EBP VpsR is activated by c-di-GMP, enhancing biofilm formation (16, 18). Similarly, the activity of the bEBP Clp can also vary based on the intracellular level of c-di-GMP in *Lysobacter enzymogenes* and *Xanthomonas campestris* (19, 20). In *E. amylovora*, the regulatory pathway by which c-di-GMP can affect T3S is not fully understood (1). The production of the two major exopolysaccharides (EPSs), amylovoran and cellulose, is also positively regulated by c-di-GMP (21, 22). *amsG*, the first gene in the amylovoran biosynthetic operon is transcriptionally regulated by c-di-GMP (4). The production of c-di-GMP is essential for biofilm formation in vitro as well as within xylem vessels in apple shoots in *E. amylovora* (9, 23).

Most studies focused on understanding the impact of c-di-GMP on critical virulence factors correlate elevated or reduced intracellular levels of c-di-GMP to downstream phenotypic outcomes (24). However, studies have not focused on understanding the regulatory impact of pGpG generation upon the degradation of c-di-GMP, specifically in terms of evaluating the role of pGpG as an independent signaling molecule. Advancements in the realm of pGpG signaling began with discovery of pGpG as an intermediate breakdown product resulting from the action of an EAL-type PDE on c-di-GMP (25). Further, Orr et al. demonstrated that Orn can degrade pGpG into GMP subunits in *V. cholerae* (6) and recently, Heo et al. discovered a PDE in *V. cholerae* which was termed PggH and could specifically degrade pGpG into GMP via DHAA1 domain activity (7). A recent review by Galperin and Chou surveyed over ∼1100 diverse bacterial genome sequences and found that while the EAL domain containing PDEs are more prevalent than HD-GYP-type PDEs, ∼250 genomes encoded more HD-GYPs than EALs while a majority, ∼580 genomes, encoded more EALs than HD-GYPs (26). Since *E. amylovora* does not encode any HD-GYPs, we hypothesized that the intermittent formation of pGpG via EAL activity could be indicative of an evolutionary adaptation whereby pGpG is used as a signaling molecule.

*E. amylovora* encodes a small repertoire of DGCs and PDEs, with five enzymatically active DGCs and three PDEs (9). In a previous study, we demonstrated that the deletion of the genes encoding for these active DGCs and PDEs in the strain Ea1189, termed Δ8, resulted in undetectable levels of c-di-GMP (9). Conversely, the previously reported strain Ea1189Δ*pdeABC,* hereafter referred to as Δ3, has high intracellular levels of c-di-GMP due to the deletion of all three active PDEs (4). These two strains, Δ8 and Δ3, provide a low and high c-di-GMP background respectively for genetic studies. To study the signaling implications of high and low pGpG/GTP levels, we heterologously expressed DGCs and PDEs containing both EAL or HD-GYP domains in various combinations in WT Ea1189 and the mutants, Δ8 and Δ3. Our results provide several lines of evidence that pGpG specifically regulates gene expression, biofilm formation, and virulence in *E. amylovora*.

## Results

### pGpG regulates global transcriptomic targets in E. amylovora

To evaluate the potential transcriptomic targets of pGpG driven regulation in *E. amylovora*, we lowered the levels of c-di-GMP in the WT strain to pGpG or GMP by overexpressing a *V. cholerae* PDE containing either an EAL (VC1086) [Ea1189/pEAL] or HD-GYP (VC1295) [Ea1189/pGYP], respectively, and performed transcriptome analysis. Filtering the results with a DESeq2 FDR p-value cutoff of 0.05 and a relative fold change of +/-1.5-fold, 72 negative and 77 positive differentially-expressed genes (DEGs) were identified in Ea1189/pEAL when compared to Ea1189/pGYP (Supplemental datasheet 1) (Fig. 1A). Among the positive DEGs (i.e. those exhibiting more gene expression with elevated pGpG), the gene ontology (GO) enrichment analysis indicated that the overrepresented functional categories included cellular localization and metabolic substrate transport including glycerol and sulfate transport genes (Figure 1A and 1B). Among the negative DEGs (i.e. those inhibited by elevated pGpG), the overrepresented functional groups included cellular localization, pathogenesis, protein transport and host interaction (Fig. 1A and 1B).

**Figure 1:**
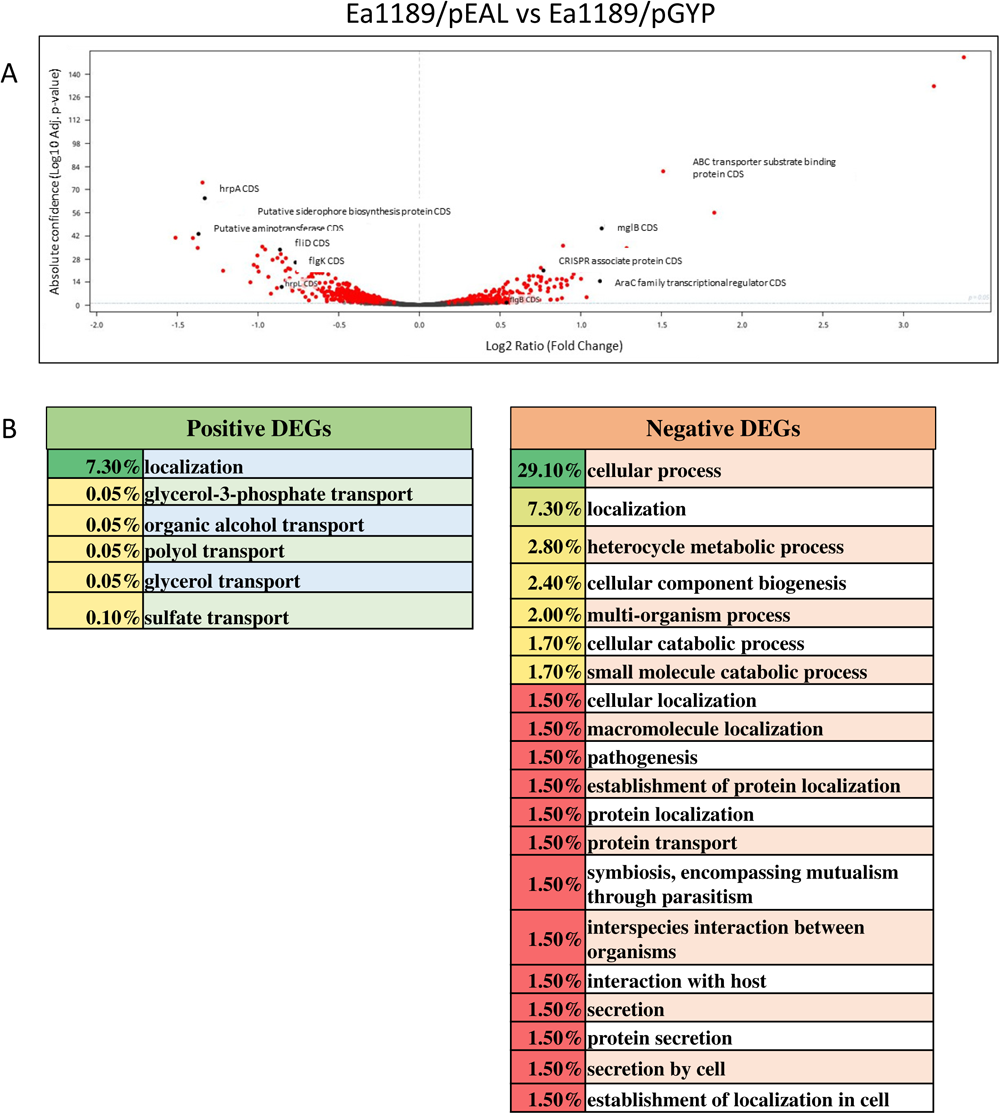
**A)** A volcano plot showing the distribution of the DEGs affected in WT/pEAL relative to WT/pGYP. The DEGs were filtered through a DESeq2 FDR cutoff of 0.05. **B)** GO enrichment analysis (FDR cutoff 0.01) highlighting the overrepresented functional groups within the positive and negative DEGs (WT/pEAL vs WT/pGYP) as a percentage of the overall DEGs.

The most positively impacted DEGs in Ea1189/EAL compared to Ea1189/HDGYP included EAM_3381and EAM_2228 which encode for *pdeC* and *pdeA* respectively, two EAL-type PDEs present in *E. amylovora* (Table 1). This result suggests that an accumulation of pGpG induces expression of the EAL-encoding PDEs, presumably to reduce its accumulation. Some of the other positive DEGs encode for hypothetical and/or putative membrane proteins (EAM_2219, EAM_2978, EAM_1000) (Table 1). Several phage response related targets were also upregulated including EAM_2114 and EAM_2113 (encode for a putative phage proteins) and phage shock protein encoding gene *pspA* (EAM_1840) (Table 1). EAM_0234 which encodes for a fimbrial protein was the main extracellular appendage related gene that was upregulated (Table 1). Two genes encoding for an AraC-family transcriptional regulator (EAM_3320 and EAM_0942) and two genes encoding a LysR-family transcriptional regulator including *nac* (EAM_2112) and *cbl* (EAM_2111) were also among the main positive DEGs (Table 1). The DEGs negatively impacted in Ea1189/pEAL compared to Ea1189/pGYP included several hypothetical proteins which were not annotated (EAM_1820, EAM_2123, EAM_2124, EAM_2127 and EAM_2128), the flagellar filament encoding gene *fliC*, type III section system genes *hrpA* and *hrpN,* and the *thiE* and *thiC* genes involved in thiamine synthesis (Table 2). Other substrate transport related genes including EAM_1726 which encodes for a TonB dependent receptor, and *sitA* and *sitB* which are involved in iron transport were also among the negative DEGs (Table 2). These results show that pGpG induces a specific transcriptome in *E. amylovora*.

**Table 1:**
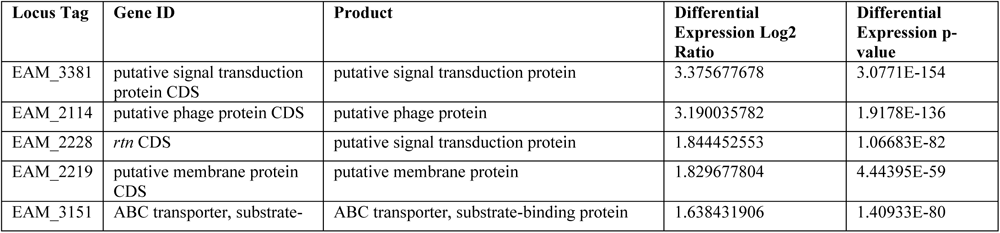
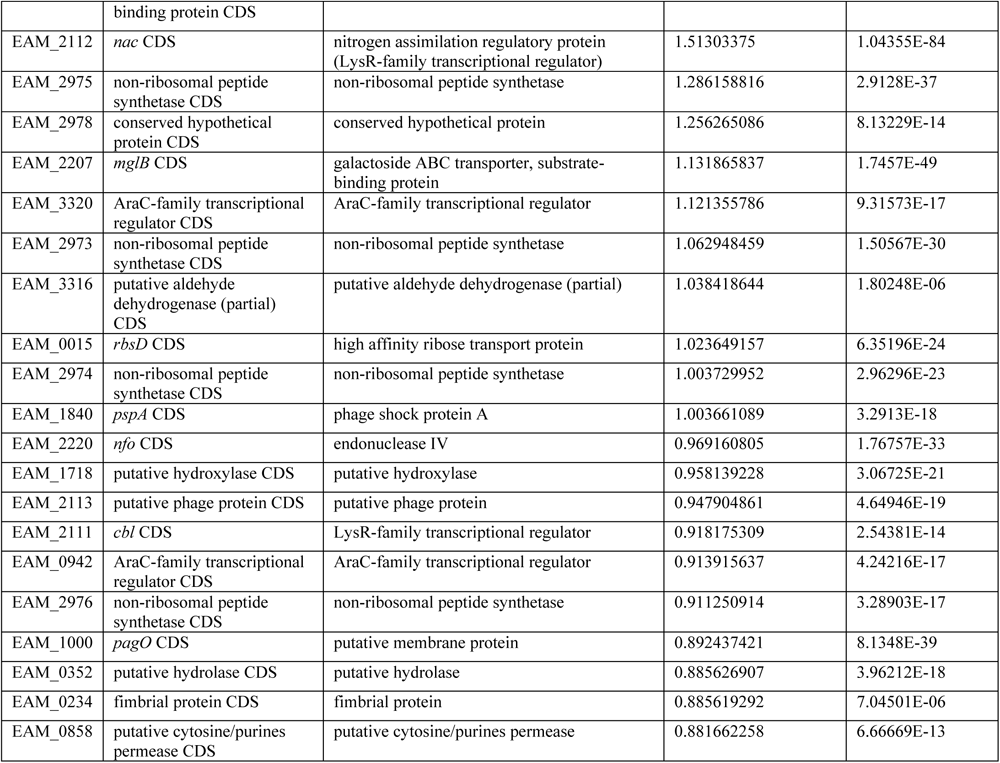
A list of the 25 most positively regulated genes in WT Ea1189/pEAL relative to WT Ea1189/pGYP.

**Table 2:** A list of the 25 most negatively regulated genes in WT Ea1189/pEAL relative to WT Ea1189/pGYP.

| Locus Tag | Gene ID | Product | Differential Expression Log2 Ratio | Differential Expression p-value |
| --- | --- | --- | --- | --- |
| EAM_2124 | conserved hypothetical protein CDS | conserved hypothetical protein | -1.509296064 | 1.05E-43 |
| EAM_2127 | conserved hypothetical protein CDS | conserved hypothetical protein | -1.401575727 | 1.71E-43 |
| EAM_2123 | conserved hypothetical protein CDS | conserved hypothetical protein | -1.371623063 | 2.06E-37 |
| EAM_2126 | putative aminotransferase CDS | putative aminotransferase | -1.36673136 | 4.48E-46 |
| EAM_2125 | putative phenylacetate-coenzyme A ligase CDS | putative phenylacetate-coenzyme A ligase | -1.341903638 | 1.32E-77 |
| EAM_2887 | *hrpA* CDS | type III secretion system protein | -1.327918425 | 5.49E-68 |
| EAM_2067 | *fliC* CDS | flagellin (FliC) | -1.218234902 | 4.25E-68 |
| EAM_2128 | conserved hypothetical protein CDS | conserved hypothetical protein | -1.214324407 | 2.63E-23 |
| EAM_1253 | *hutU* CDS | urocanate hydratase | -1.144439758 | 3.13E-48 |
| EAM_0240 | *thiC* CDS | thiamine biosynthesis protein | -1.123480245 | 2.45E-43 |
| EAM_0359 | putative siderophore biosynthesis protein CDS | putative siderophore biosynthesis protein | -1.096472308 | 3.73E-55 |
| EAM_2877 | *hrpN* CDS | harpin | -1.07000488 | 6.01E-47 |
| EAM_0239 | *thiE* CDS | thiamine-phosphate pyrophosphorylase | -1.04514606 | 5.06E-16 |
| EAM_0360 | putative siderophore biosynthesis protein CDS | putative siderophore biosynthesis protein | -1.036484327 | 2.78E-60 |
| EAM_0361 | putative decarboxylase CDS | putative decarboxylase | -1.02421314 | 6.10E-27 |
| EAM_2129 | putative ornithine cyclodeaminase CDS | putative ornithine cyclodeaminase | -1.002578394 | 1.02E-25 |
| EAM_1726 | TonB-dependent receptor CDS | TonB-dependent receptor | -1.001690728 | 9.18E-33 |
| EAM_2383 | *flxA* CDS | conserved hypothetical protein | -0.994002784 | 9.14E-23 |
| EAM_2087 | *sitA* CDS | iron ABC transporter, substrate-binding protein | -0.986485239 | 9.76E-67 |
| EAM_2088 | *sitB* CDS | iron ABC transporter, ATP-binding protein | -0.971214483 | 3.30E-38 |
| EAM_1259 | *hutG* CDS | N-formylglutamate amidohydrolase | -0.953742463 | 1.79E-36 |
| EAM_1254 | *hutH* CDS | histidine ammonia-lyase | -0.93363205 | 1.90E-19 |
| EAM_1820 | hypothetical protein CDS | hypothetical protein | -0.920824941 | 1.94E-45 |
| EAM_0873 | ABC transporter, permease protein CDS | ABC transporter, permease protein | -0.917911247 | 9.08E-09 |

### Generation of distinct intracellular concentrations of c-di-GMP and pGpG

In order to study the dynamics of pGpG driven signaling in *E. amylovora* we had to distinguish the impact of pGpG from the loss of c-di-GMP. To accomplish this, we used five strains that compromised varying intracellular concentrations of c-di-GMP or pGpG as measured by LC-MS/MS. WT Ea1189 was used as the primary control strain, and it exhibited low but detectable levels of both c-di-GMP and pGpG (4). Δ3, lacking all three PDE-encoding genes (*pdeABC*) is a strain with no c-di-GMP degradation activity and thus exhibited high c-di-GMP and no detectable pGpG (Fig. 2A, B). Overexpression of the pEAL or pGYP plasmids described above decreased c-di-GMP concentrations of Δ3 to the WT level (Fig. 2A), but only in the case of pEAL did pGpG accumulate (Fig. 2B). No pGpG is detectable in Δ3/pGYP (Fig. 2B). Δ8, which is a deletion mutant lacking all five active DGC (*edcA-E*) and three PDE (*pdeA-C*) encoding genes (9), has no detectable c-di-GMP or pGpG (Fig. 2A, B). Thus, this set of strains yielded various intracellular states of c-di-GMP and pGpG that could be further analyzed for phenotypic output to assess any potential signaling impact of pGpG in *E. amylovora*. We also hypothesized that the intracellular concentration of GTP could be altered in these five strains.

**Figure 2:**
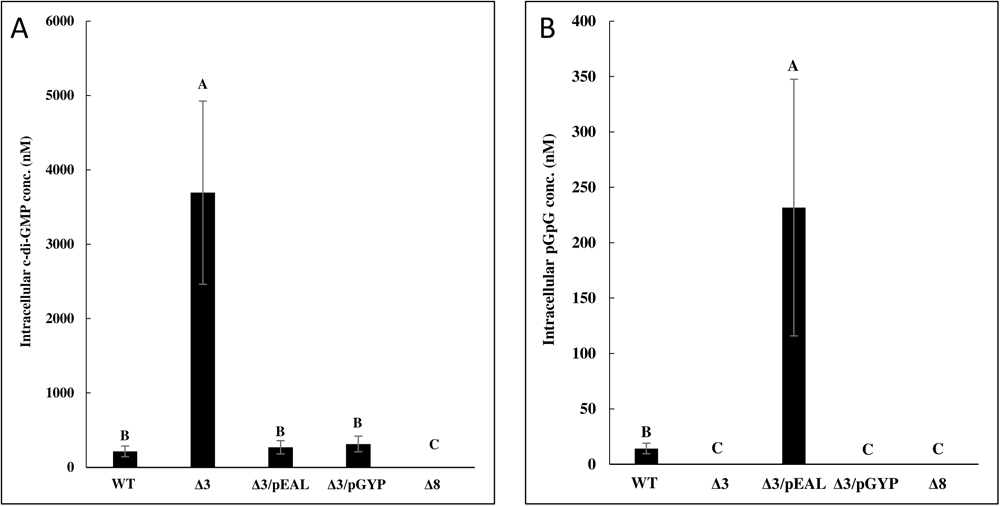
Intracellular levels of **A)** c-di-GMP and **B)** pGpG in WT Ea1189, Δ8 and Δ3 expressing pEAL and pGYP in different combinations. Error bars represent standard error of the means and significance letters denote statistically significant (p-value 0.05) differences determined via Tukey’s HSD.

GTP levels were highest in WT Ea1189 and this was significantly higher than the four other strains (Fig. S1). Some significant differences in GTP concentrations were observed between Δ3, Δ3/pEAL, Δ3/pGYP, and Δ8, but the overall differences were marginal relative to the changes in c-di-GMP and pGpG (Fig. S1).

### The degradation of c-di-GMP into pGpG or GMP can have differential impacts on amylovoran production and biofilm formation

The first phenotype we examined was amylovoran production. Amylovoran is an extracellular polysaccharide that is important for biofilm formation and virulence and the biosynthesis of amylovoran is induced by c-di-GMP (4, 9, 27). Mutant Δ3, which has high c-di-GMP and no pGpG, exhibited elevated amylovoran production compared with the WT strain, supporting that c-di-GMP can induce its production. Δ3*/*pEAL also exhibited significantly elevated levels of amylovoran when compared to WT Ea1189, even though this strain has low c-di-GMP but elevated pGpG (Fig. 3A). Surprisingly, Δ3*/*pGYP showed the highest level of amylovoran production at levels that were significantly higher than Δ3 and Δ3*/*pEAL. The amount of amylovoran produced by Ea1189Δ8 was equivalent to WT (Fig. 3A). *In vitro* biofilm formation of these five strains closely mimicked amylovoran production (Fig. 3B). These results suggest that any alteration in the ratio of c-di-GMP/pGpG induces amylovoran and biofilms. Unlike biofilms, flagellar motility was relatively unchanged for all the strains evaluated including WT Ea1189, Δ3, Δ3*/*pEAL, Δ3*/*pGYP and Δ8 (Fig. 3C).

**Figure 3:**
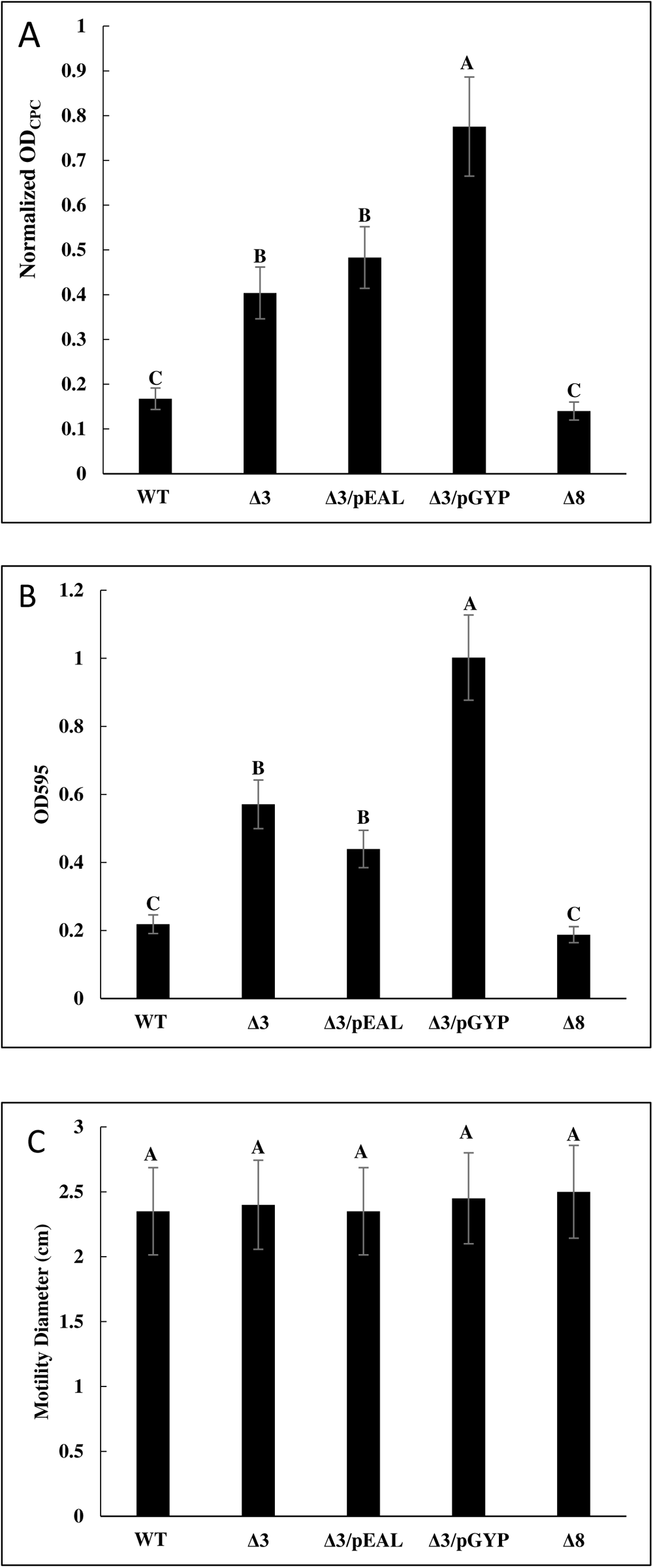
Relative levels of **A)** amylovoran production, **B)** biofilm formation and **C)** flagellar motility measured for WT Ea1189, Δ8 and Δ3 expressing pEAL and pGYP in different combinations. Error bars represent standard error of the means and significance letters denote statistically significant (p-value 0.05) differences determined via Tukey’s HSD.

### Impact of c-di-GMP and pGpG on expression of hrpL and hrpS

We next turned our analysis to the impact of pGpG on virulence gene. A major virulence factor of *E. amylovora* is the Type III secretion system, which is regulated by a complex array of transcription factors ultimately impinging on expression of the alternative sigma factor HrpL. As mentioned in the introduction, HrpS is an EBP-family transcription factor that induces expression of *hrpL.* We therefore used q-RT-PCR expression to assess the impact of c-di-GMP and pGpG in the five strains we have examined thus far. Transcript levels of *hrpS* were similar in WT, Δ3, Δ3/pGYP, and Δ8 (Fig. 4A). Interestingly, Δ3/pEAL had *hrpS* transcript levels that were significantly increased 5-fold increase compared to all other evaluated strains (Fig. 4A).

**Figure 4:**
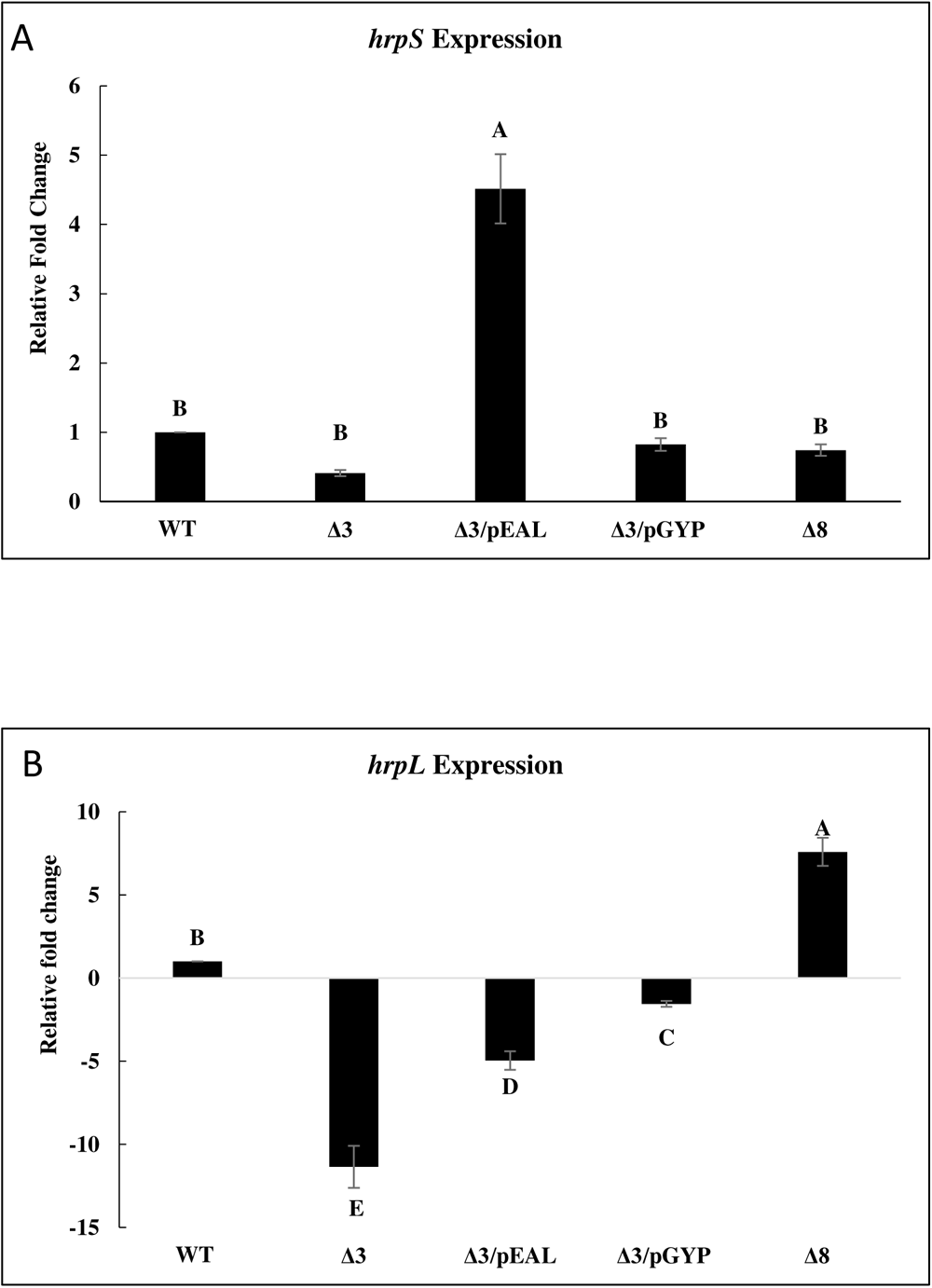
Fold change in **A**) *hrp*S and **B)** *hrpL* transcript levels in WT Ea1189, Δ8 and Δ3 expressing pEAL and pGYP in different combinations. Error bars represent standard error of the means and significance letters denote statistically significant (p-value 0.05) differences determined via Tukey’s HSD.

The transcript levels of *hrpL*, as measured by q-RT-PCR, however, did not correlate with *hrpS* expression. Expression of *hrpL* was significantly reduced in Δ3 relative to WT Ea1189, supporting previous results that c-di-GMP inhibited *hrpL* expression (Fig. 4B) (4, 9). Δ3/pEAL and Δ3/pGYP showed partial inhibition of *hrpL* expression relative to Δ3, although expression remained below WT levels, suggesting that an improper ratio of c-di-GMP/pGpG negatively impacted *hrpL* (Fig. 4B). Mutant Δ8 exhibited the highest transcript levels of *hrpL*, which were significantly different from all other strains, showing that the lack of any c-di-GMP signaling molecules led to the greatest expression (Fig. 4B).

These results show that *hrpS* expression is induced by pGpG concentrations while *hrpL* is repressed by c-di-GMP. This finding is supported by plotting the expression of *hrpS* and *hrpL* in the five examined strains to their relative concentrations of pGpG or c-di-GMP, respectively (Fig. S2). These data can be modeled with a polynomial regression fit for each gene (*hrpS/*pGpG, order 2 [Fig. S2A]; *hrpL*/c-di-GMP, order 3 [Fig. S2B]) with high confidence (R^2^ > 0.99 for both cases). Whether such relationships are causative or merely correlative requires further mechanistic insight into the transcription machinery that controls expression of *hrpS* and *hrpL* in response to pGpG and c-di-GMP.

### Model dependent impacts of c-di-GMP and pGpG on E. amylovora plant infection

Both biofilm formation and expression of the type III secretion system, mediated by HrpS and HrpL, are required for plant infection by *E. amylovora*. We assessed the virulence levels of the five strains described above with varying intracellular concentrations of c-di-GMP and pGpG in two infection models. First, we examined tissue necrosis in immature pears. This necrosis is primarily driven by the Type III secretion system and is dependent on amylovoran production during the infection process; however, biofilm formation is not critical during the progress of infection in fruit tissue (1, 27, 28). The results generally reflected *hrpL* expression levels with the WT causing large necrotic lesions, the Δ3 exhibiting the lowest lesion size, and the Δ3/pEAL and Δ3/pGYP exhibiting intermediate virulence (Fig. 5A). In this assay, however, necrosis caused by the Δ8 mutant was equivalent to the WT strain, which we interpret to mean that the WT expresses sufficient *hrpL* (and thereby Type III secretion) to cause disease in this model. Like *hrpL* expression, these results are generally correlated with intracellular c-di-GMP concentrations.

**Figure 5:**
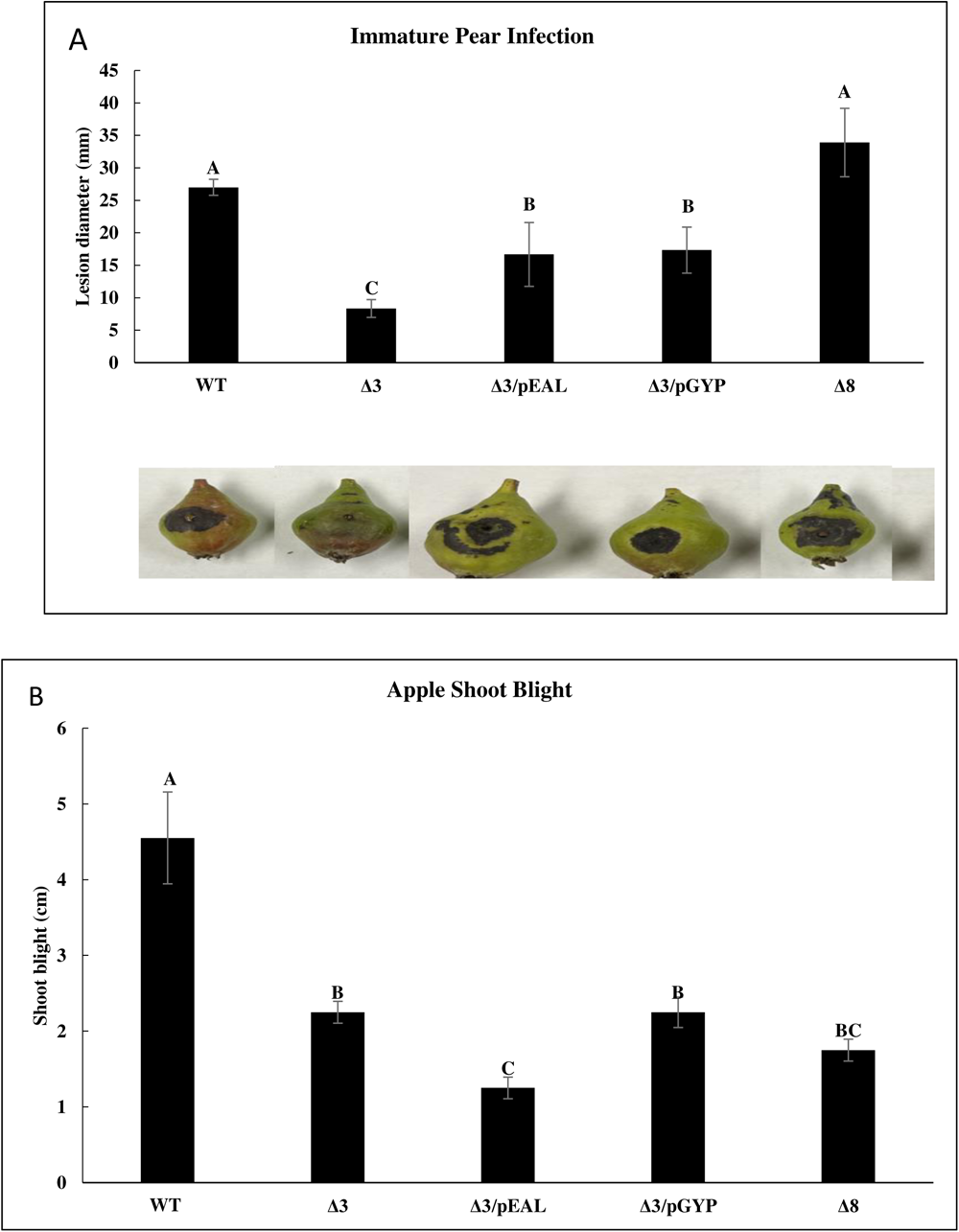
Virulence levels of WT Ea1189, Δ8 and Δ3 expressing pEAL and pGYP in different combinations in **A)** immature pears at 5dpi and **B)** apple shoots at 8dpi. Error bars represent standard error of the means and significance letters denote statistically significant (p-value 0.05) differences determined via Tukey’s HSD.

The second infection model we examined was infection of apple shoots. This is a more complex model in which *E. amylovora* utilizes Type III secretion in the apoplast tissue to initiate infection, but then migrates to the xylem vessels and generates biofilms to systemically colonize the host (2, 29, 30). Amylovoran production is essential for initial infection as well as biofilm formation (9, 27). Infection of the five strains under examination in this model exhibited a substantially different outcome than pear infection. In this case, shoot blight-based necrosis levels in apple shoots were significantly reduced in all four strains compared to WT Ea1189 (Fig. 5B). Δ3/pEAL, which had the highest concentrations of pGpG, exhibited the lowest levels of shoot blight, which were significantly reduced compared to all other strains (Fig. 5B). The results of this model show that both c-di-GMP and pGpG are necessary for optimal *E. amylovora* infection as the Δ8 strain shows reduced shoot blight lesions. Moreover, these signals must be in proper balance as found in the WT strain as either high c-di-GMP or high pGpG leads to reduced infection.

## Discussion

Although pGpG has been implicated as an inhibitor of EAL enzymatic activity (7), its role as a signaling molecule is poorly understood. Here we present several lines of evidence that the intracellular concentration of pGpG, in addition to c-di-GMP, is important for *E. amylovora* pathogenesis. First, we identified 149 genes that are differentially expressed in *E. amylovora* in a strain that degrades c-di-GMP into pGpG via an EAL PDE versus a strain that degrades c-di-GMP into 2 GMPs via an HD-GYP PDE. Secondly, we found that amylovoran production and subsequent biofilm formation is not solely dependent on c-di-GMP concentrations, but rather the balance of c-di-GMP to pGpG. Δ3/pEAL and Δ3/pGYP, which have WT concentrations of c-di-GMP, but altered pGpG concentrations, exhibit elevated biofilm formation. Such a result is inconsistent with a model that relies on only intracellular c-di-GMP. Third, we showed that transcription of *hrpS* is strongly correlated with the intracellular concentration of pGpG and not c-di-GMP. Finally, infection of apple shoots, which is a complex model that requires T3S, biofilm formation, and motility, is optimal in the WT strain that balances the concentrations of c-di-GMP and pGpG versus all other strains that exhibit altered c-di-GMP and pGpG ratios. Although in this work we do not define the molecular mechanism by which c-di-GMP and pGpG are sensed and alter phenotypic behaviors, these findings provide evidence that the intracellular concentration of pGpG is important for disease progression in *E. amylovora*.

Our transcriptomic study aimed at evaluating the global impact of rising levels of pGpG in *E. amylovora* revealed several DEGs involved in metabolism, virulence, extracellular appendage generation and general transcriptional regulation. Transcriptomic studies so far have mostly focused on understanding the global impact of c-di-GMP levels within pathogenic bacteria (31–33). Our previous study in *E. amylovora* highlighted that the presence or absence of c-di-GMP during the initial stages of biofilm formation can have global transcriptomic implications, especially on several metabolic processes (9). In our study, several genes encoding transcriptional regulators of the LysR and AraC family were upregulated when higher levels of pGpG were present in the cell. AraC family regulatory proteins have been demonstrated to be able to bind to c-di-GMP, but to our knowledge, pGpG has not been evaluated as a potential ligand or regulatory factor affecting AraC family protein transcription and/or regulation (34, 35). Targets that were downregulated by higher cellular pGpG levels included *hrpA* and *hrpN* which encode for components of the T3S in *E. amylovora* (36). In *E. amylovora*, the expression of T3S genes is negatively regulated by c-di-GMP (4). Inhibition of T3S by pGpG could explain the reduced virulence of Δ3/pEAL in the apple shoot infection model.

An evaluation of the levels of *hrpL* expression indicated a strong negative correlation between c-di-GMP levels and *hrpL* transcripts. Alternatively, elevated pGpG levels induced the expression of *hrpS*. Thus, *hrpL* and *hrpS* expression respond differently to intracellular levels of c-di-GMP and pGpG. HrpL functions as an alternate sigma factor in *E. amylovora* and the activity of HrpL can positively regulate the transcription of other *hrp* genes including *hrpA* and *hrpN* (10, 36). HrpS is an enhancer binding protein that interacts with the alternative sigma factor 54 (RpoN) and its modulation protein YhbH to regulate *hrpL* transcription in *E. amylovora* (12). The negative correlation between c-di-GMP levels and *hrpL* expression has been studied in many phytopathological systems including *E. amylovora* (3), *D. dadantii* (37) and *P. syringae* (31). In *E. amylovora*, the overexpression of the DGC encoding gene *edcC* resulted in an elevation in *hrpS* promoter activity (3). In *D. dadantii,* the deletion of two different PDE encoding genes *ecpB* and *ecpC* did not change the promoter activity of *hrpS* (38). To our knowledge, no studies have linked pGpG to *hrpS* expression. In *E. amylovora*, our data suggests a strong positive correlation between intracellular pGpG levels and *hrpS* expression. However, the molecular mechanism underlying this effect needs further investigation.

T3S expression and function are critical factors that determine the virulence outcomes on immature pear fruit (28). Corresponding to the lower than WT levels of *hrpL* expression observed in Δ3/pEAL and Δ3/pGYP, virulence in immature pear was reduced in these strains relative to WT Ea1189. The reduced virulence levels in Ea1189Δ3 have been previously attributed to the significantly elevated c-di-GMP levels observed in this strain, which can lead to reduced T3S expression and DspE translocation (4). However, no distinct differences in pear infections were observed when the c-di-GMP in Δ3 was degraded by an EAL or HD-GYP-type PDE, suggesting c-di-GMP is the primary driver of virulence in this model.

While infecting apple shoots, *E. amylovora* must employ T3S in the leaf apoplast as a means to suppress host defense responses and biofilm formation in the xylem vessels to mediate systemic spread (1). The degradation of c-di-GMP in Δ3 by the action of an EAL-type PDE resulted in a decline in shoot blight levels, which was not the case in Δ3/pGYP. C-di-GMP is essential for biofilm formation in xylem vessels during the *E. amylovora* infection process (2, 9). Due to the positive regulatory impact on *hrpL* expression arising from pGpG and GMP generation through the hydrolysis of c-di-GMP, the reduction of shoot blight levels in Δ3/pEAL could be attributed to a reduction in biofilm formation and systemic spread by these strains. Similar to the trend observed with biofilm formation in vitro, the proportional balance of c-di-GMP and pGpG seems to be critical for successful shoot blight infection, and any changes in this ratio resulting from the addition/deletion of DGCs and/or PDEs can negatively affect virulence in apple shoots. There are some experimental limitations that could impact our observations. Overexpression was induced throughout the in vitro biofilm and amylovoran assays, while for the in vivo experiments the strains were induced prior their inoculation in planta. Also, were are currently unable to quantify c-di-GMP, pGpG and GTP levels within pathogen cells during disease progression (1, 24, 39).

EAL-type PDE activity on c-di-GMP resulting in pGpG formation has been studied in several systems including *D. dadantii*, *Xanthomonas oryzae* and *P. aeruginosa* (38, 40–42). These studies have highlighted specific PDEs EcpB and EcpC in *D. dadantii* that can enzymatically hydrolyze c-di-GMP into pGpG and can positively regulate pectate lyase production impact virulence in the host (38). While *X. oryzae* does encode for HD-GYP-type PDEs, PdeR, an EAL-type PDE is a strong positive regulator of virulence and EPS production (40, 41). RocR an EAL-type PDE in *P. aeruginosa* was shown to be able to cleave c-di-GMP into pGpG and thus contribute to biofilm dispersal (42). One common factor among these studies was the correlation of c-di-GMP degradation to virulence related phenotypes and not a direct evaluation of pGpG as a signaling factor. Our results suggest that both c-di-GMP and pGpG are important signals regulating disease in *E. amylovora*.

## Materials and Methods

### Bacterial strains, plasmids and growth conditions

The bacterial strains, plasmids and their relevant characteristics are listed in Table 3.

**Table 3:** Strains, plasmids and their relevant characteristics.

| Bacterial strain or plasmid | Relevant characteristics | Reference |
| --- | --- | --- |
| *E. amylovora* strains |  |  |
| Ea1189 | Wild type |  |
| Δ3 | *pdeA*, *pdeB* and *pdeC* deletion mutant in Ea1189 | (4) |
| Δ8 | *edcA-E* and *pdeA-C* deletion mutant in Ea1189 | (9) |
| Plasmids |  |  |
| pEVS143 | Broad-host range, IPTG inducible (Ptac) cloning vector; Km<sup>R</sup> | (57) |
| pACYCDuet-1 | Expression vector containing two MCS: p15A ori; Cm<sup>R</sup> | Novagen, Germany |
| pEAL | *V. cholerae* gene VC1096 in pEVS143; Km<sup>R</sup> | (58) |
| pGYP | *V. cholerae* gene VC1295 in pEVS143; Km<sup>R</sup> | (59) |

Strains were grown in lysogeny broth (LB) at 28°C unless otherwise noted with the appropriate antibiotics at the following concentrations: ampicillin (Ap; 100 µg/ml), chloramphenicol (Cm; 10 µg/ml) and kanamycin (Km; 100 µg/ml). Strains harboring an overexpression vector were induced with 1mM iso-propyl-β-D-thiogalactopyranoside (IPTG).

### Transcriptomics, bioinformatics, q-RT-PCR and data analysis

*E. amylovora* WT Ea1189 harboring pEAL or pGYP were grown overnight, sub-cultured until they reached the mid-log phase harvested and resuspended in Hrp-inducing minimal medium (HRP-MM) (43) for 6h with IPTG induction. Cells were collected and processed for RNA-Seq analysis according to a previously described workflow (9). RNA extraction from the samples was conducted according to the protocol by Rivas et al. (44). Washing the samples with 0.1% N-lauryl sarcosine sodium salt preceded heat-based lysis using the lysis buffer (1% SDS in 10mM EDTA and 50mM sodium acetate, pH 5.1). The RNA clean and concentrator kit (Zymo Research, CA, USA) along with the TURBO DNA-free kit (Thermo Fisher Scientific, MA, USA) were used to treat the samples for DNA contamination and to concentrate the RNA. The Agilent 4200 TapeStation (Agilent Technologies, CA, USA) was used to analyze the quality and RIN-values of the RNA. The RNA was treated with the QIAseq FastSelect 5S/16S/23S rRNA removal kit prior to library prep with the TruSeq Stranded Total RNA Library Prep Kit (Illumina, CA, USA). The Illumina HISeq 4000 was used for sequencing at 50bp single-end reads.

Raw reads were processed for barcode filtering using Trimmomatic v 0.36 (singleendcri-teria:ILLUMINACLIP:TruSeq3-SE:2:30:10LEADING:3TRAILING:3SLIDINGWIN-DOW:4:15) (45). The *E. amylovora* Ea1189 genome downloaded from NCBI was used as a reference to map trimmed sequences with Bowtie v 2.1.1, followed by gene expression determination with HTSeq v 0.11.2 (46–48). DESeq2 v 3.12 was used for differential gene analysis with an FDR cutoff of 0.05 (p-value) and a minimum fold change of 1.5-fold (log2) (49). Cytoscape/BiNGO (FDR cutoff 0.01) was used for gene ontology enrichment analysis and Geneious software was used for graphical data analysis (50, 51). Three biological replicates were included in the experiment.

q-RT-PCR was used to quantify *hrpL* and *hrpS* gene expression as previously described (4). RNA extraction and cleanup was conducted as per the process used for RNA-Seq. cDNA was synthesized from the RNA using the High-Capacity Reverse Transcriptase kit (Applied Biosystems, CA, USA). The SYBR green PCR master mix (Applied Biosystems, CA, USA) was used to conduct quantitative PCR experiments. The delta C_T_ method was used to determine changes in levels of gene expression (52). Three biological replicates were included in the experiments.

For statistical analysis of data acquired in all phenotypic and q-RT-PCR based assays, the Tukey’s honestly significant difference (HSD) was used in the JMP™ statistical software platform.

### Quantification of c-di-GMP, pGpG, and GTP

Intracellular levels of c-di-GMP and pGpG were quantified by a previously described protocol using ultra performance liquid chromatography and tandem mass spectrometry (UPLC-MS/MS) (3, 6). Strains were grown overnight in LB, sub-cultured and induced for 6h with IPTG as appropriate. Cells were collected, lysed (40% acetonitrile and 40% methanol) and analyzed against synthesized standard of c-di-GMP (Axxora Life Sciences Inc., CA, USA), pGpG (Jena Bioscience, Germany), and GTP (Jena Bioscience, Germany) using the Quattro-Premier XE instrument (c-di-GMP and pGpG) (4, 6) or Acquity UPLC Premier BEH as previously described (53). Three biological replicates were included in the experiments.

### Quantification of amylovoran production

Amylovoran production was measured in the strains using a previously described protocol (4). Strains were grown in MBMA medium (54) amended with antibiotics and IPTG as appropriate for 48h at 28°C. Relative levels of amylovoran production were quantified in the supernatants of grown cultures using cetylpyridinium chloride CPC (50 mg/ml). The ratio of OD_600_ of the CPC precipitate to the OD_600_ of the initial cell culture density was used as a relative measure of amylovoran production. Three biological replicates were included in the experiments.

### Quantification of biofilm formation

Biofilm formation was quantified using a previously described protocol (55). Strains were grown overnight in LB followed by OD_600_ normalization and resuspension in 0.5X LB amended with antibiotics and IPTG as appropriate for 48h along with three polypropylene beads (diameter 7mm). A 0.1% crystal violet solution was used to stain the beads for 1h and 1 ml of elution buffer (40% methanol and 10% glacial acetic acid). The OD_595_ was measured of the eluted solution and used as a relative representation of biofilm formation by the strains. Three biological replicates were included in the experiments.

### Flagellar motility

The swimming motility assay was used to measure flagellar motility within the strains as previously described (55). Strains were grown overnight and their OD_600_ was normalized. A sterile 10 µL pipet tip was dipped in the inoculum and stabbed onto a motility agar plate (10 g tryptone, 5 g NaCl and 0.3% agar/liter) amended with IPTG as appropriate. The plates were incubated at 28°C for 24h. Relative flagellar motility levels were quantified by analyzing the cellular spread using ImageJ (56). Three biological replicates were included in the experiments.

### Virulence assays

For both the apple shoot blight and immature pear infection assays, the same strain culturing conditions were used. Strains were grown overnight in LB followed by sub-culturing and induction with IPTG as appropriate for 6h. The strains were then used for each of the specified virulence assays.

The immature pear fruit infection assay was performed as previously described (4). The pears were stab-inoculated at a concentration of 10^4^ CFU/ml. The pears were incubated under conditions of high-humidity for 5 days at 28°C. Infection levels were determined by measuring the diameter of the necrotic lesion developing around the wound site. The shoot blight assay was conducted as previously described (4). Newly growing shoot tips on apple trees (*Malus x domestica* cv. Gala) were infected by cutting between the peripheral veins using scissors dipped in cell cultures normalized to an OD_600_ of 0.8. Shoot blight levels were quantified by measuring necrosis along the shoot developing from the site of inoculation. Three biological replicates were included in the experiments.

## Acknowledgements

This project was funded by NIH grants AI144395 to G.W.S. and C.M.W. and NIH grants GM139537 and AI158433 to C.M.W.

## Data availability

RNA Sequencing reads and data files are available on NCBI under the BioProject accession number PRJNA1042810.

**Supplemental Datasheet 1**: The complete DEG lists for the RNA-Seq conducted in this study comparing Ea1189/pEAL vs Ea1189/pGYP.

**Figure S1:** Intracellular levels of GTP in WT Ea1189, Δ8 and Δ3 expressing pEAL and pGYP in different combinations. Error bars represent standard error of the means and significance letters denote statistically significant (p-value 0.05) differences determined via Tukey’s HSD.

**Figure S2:** Polynomial regression fit analysis for **A)** *hrpS* expression and pGpG, (order 2 regression) and **B)** *hrpL* expression and c-di-GMP (order 3 regression). The trendline equation and R^2^ values for each of the models are listed on the graph.

